# SARS-CoV-2 Protein Nsp2 Stimulates Translation Under Normal and Hypoxic Conditions

**DOI:** 10.1101/2022.09.13.507829

**Authors:** Nadejda Korneeva, Md Imtiaz Khalil, Ishita Ghosh, Ruping Fan, Thomas Arnold, Arrigo De Benedetti

## Abstract

When viruses like SARS-CoV-2 infect cells, they reprogram the repertoire of cellular and viral transcripts that are being translated to optimize their strategy of replication, often targeting host translation initiation factors, particularly eIF4F complex consisting of eIF4E, eIF4G and eIF4A. A proteomic analysis of SARS-CoV-2/human proteins interaction revealed viral Nsp2 and initiation factor eIF4E2, but a role of Nsp2 in regulating translation is still controversial. HEK293T cells stably expressing Nsp2 were tested for protein synthesis rates of synthetic and endogenous mRNAs known to be translated via cap- or IRES-dependent mechanism under normal and hypoxic conditions. Both cap- and IRES-dependent translation were increased in Nsp2-expressing cells under normal and hypoxic conditions, especially mRNAs that require high levels of eIF4F. This could be exploited by the virus to maintain high translation rates of both viral and cellular proteins, particularly in hypoxic conditions as may arise in SARS-CoV-2 patients with poor lung functioning.

## INTRODUCTION

The rapid spread of the Covid-19 pandemic since the beginning of 2020 can only be described as cataclysmic. While it caused immense worldwide hardship and suffering, it also caused a paradigm shift in the way scientists were called into action to respond to a previously never encountered urgent societal need. The scientific community has joined efforts to collect knowledge and push the therapeutic response to Severe acute respiratory syndrome coronavirus 2 (SARS-CoV-2) infection. New collaborations are being born every day and scientific data are shared widely as soon as they become available. From the super-rapid sequencing of new variants and structural determination of the viral genome organization and immediate comparison to SARS-CoV and MERS-CoV (1), to the overwhelming recent feat of identifying the complete interactome of cellular and virally-encoded proteins by an international consortium spearheaded by the Krogan lab at unprecedented speed (2), which has led to better understanding of the virus.

SARS-CoV-2 is a betacoronavirus that has large positive single-stranded mRNA genomes containing cap-structure (m^7^GpppN-) and (polyA)-tail at their 5’ and 3’-ends, respectively (3). Like other positive-strand RNA viruses, after entering cell, SARS-CoV-2 RNA is transported to endoplasmic reticulum (ER) to be translated by a host translation machinery. The 5’-two-thirds of the ~30kb SARS-CoV-2 genome has two overlapping open reading frames (ORF), the ORF1a and ORF1b (4). Polyprotein (pp) 1a is synthesized from the ORF1a. A second polyprotein, pp1b, that is encoded by both ORF1a and ORF1b requires a −1 ribosomal frameshift during elongation step of translation. Newly synthesized polyproteins are co-translationally and post-translationally processed into the individual non-structural proteins (Nsp). They together comprise 15-16 Nsps including Nsp1 to Nsp10 and Nsp12 to Nsp16. These Nsps are involved in assembly of viral replication complex and transcription of genomic mRNAs (5). The ORFs located in the 3’-end of the genomic RNA encode structural (S, E, M, and N) and accessory (3a, 3b, 6-10) proteins that are synthesized from sub-genomic mRNAs generated by discontinued transcription. Similarly to genomic RNA, all sub-genomic transcripts have 5’-cap and 3’-poly(A)-tail. In addition, all sub-genomic SARS-CoV-2 RNAs have a common 5’-leader sequence (4).

The presence of 5’-cap on genomic and sub-genomic SARS-CoV-2 mRNAs indicates ability of viral mRNAs to utilize a host translation machinery to synthesize viral proteins. Initiation of translation is a highly regulated step of protein synthesis (6). It involves sequential assembly 43S, 48S, and 80S initiation complexes. The binding of mRNA to the 43S to form 48S is regulated by the eIF4F protein complex that consist of eIF4E, eIF4G, and eIF4A. The eIF4F complex facilitates 5’-cap recognition of mRNA via eIF4E interaction with the cap, unwind mRNA secondary structure in the 5’-UTR via helicase activity of eIF4A, and loads mRNA onto 43S pre-initiation complex via the eIF4G protein-binding activities. Cap-dependent translation is a major mechanism of initiation of translation in eukaryotic cells. However, under stress, viral infection or metabolic changes, translation of some cellular mRNA to be initiated at the internal ribosome entry site (IRES) located in their 5’-UTR (7). Switch from the cap-dependent to cap-independent translation allows cells to synthesize proteins necessary for their adaptation to stress and survival. The level of eIF4E available to form eIF4F complex determines whether mRNA will be initiated via the cap- or IRES-dependent mechanism. Sequestration of eIF4E from the eIF4F complex is regulated by its binding to the repressor protein 4EBP, which is, in turn, regulated by the PI3K/Akt/mTOR signaling cascade (8). Upon phosphorylation of 4EBP by mTOR during mitogenic and nutrient sufficiency, it dissociates from eIF4E, thus allowing formation of eIF4F complex and cap-dependent translation (9).

To compete with cellular mRNAs, coronaviruses use several strategies (4). SARS Nsp1 protein was shown to shut off host protein synthesis by targeting initiation step of protein synthesis (10), (11), (12). It was shown that SARS-CoV-2 Nsp1 interacts with the 37-nt region of 40S that is adjacent to the mRNA entry channel (13), (14), (15). In addition to 40S, Nsp1 binds untranslated 80S ribosomes, thus depleting ribosome from the translating pool (15). These interactions block 40S scanning along host mRNAs and disrupt tRNA loading to 80S, all leading to inhibition of global translation (13), (15). Interestingly, SARS Nsp1 was shown to inhibit not only cap-dependent translation but also translation of some mRNAs containing IRES in their 5’-UTR (16), (17), (12). At the same time, translation of the viral sub-genomic mRNAs is not inhibited by Nsp1 due to the presence of the viral leader sequence in their 5’-ends (13). Moreover, it was demonstrated that a reporter containing full-length 5’-UTR of genomic SARS-CoV-2 mRNA translated more efficiently in the presence of Nsp1 (15). In addition, upon binding, SARS Nsp1 was shown to induce endonucleolytic cleavage of host mRNAs near the 5’UTR, targeting them for degradation (16), (17). At the same time, viral mRNAs are protected from cleavage due to the presence of a 5’-end leader sequence (17). In order to suppress translation, other SARS proteins interact with host translation factors (4). It was reported that SARS-CoV spike protein S inhibits host translation via interaction with eIF3 subunit f (18). Expression of SARS-CoV N protein was shown to induce the aggregation of elongation factor eEF1α *in vivo* and in *in vitro* experiments and inhibited host translation (19). SARS-CoV protein 7a was shown to not only inhibit host translation but also to induce cell apoptosis (20).

Recently, a proteomic analysis identified potential interaction between SARS-CoV-2 protein Nsp2 and host translation initiation factor eIF4E2 (2), (21), (22). Among coronaviruses, Nsp2 is a very conserved protein. However, the SARS-CoV-2 Nsp2 protein appears to be still under evolutionary pressure (23). Beside its possible role in mitochondrial (24), (25), (21) and endosomal biogenesis (26), the function of SARS-CoV Nsp2 is still unknown. It was demonstrated that while SARS-CoV Nsp2 is dispensable for viral replication in cell culture, its deletion affected viral growth and RNA synthesis (27). eIF4E2 or 4EHP (4E-homologous protein), a paralog of the cap binding protein eIF4E, shares 28% identity with mammalian eIF4E (28), (29), (30). It was detected in various organisms and was shown to regulate proper embryonic development in *Drosophila* (31), (32), and in mammals (33), to be essential for murine germ cell development (34) and in miRNA-mediated silencing in *C. elegans* (35), (36). eIF4E2 has lower expression level in mammalian cell lines compared to eIF4E and reportedly a weaker affinity for the 5’-cap mRNA structure (37), (38), (29). *Schizosaccharomyces pombe* eIF4E2 was shown to bind eIF4G more than 100-fold more weakly than eIF4E1 *in vitro* (39). Mammalian eIF4E2 was shown to bind 4EBP1 in cells but this interaction does not seem to respond to the traditional mTOR pathway of protein synthesis stimulation, suggesting a weaker binding compared to eIF4E to 4EBP1 (40), (38). The proposed role of eIF4E2 as a translational inhibitor is based on its ability to form specific protein complexes on both 5’ and 3’-UTRs of a target mRNA that interferes with the recognition of the 5’-cap mRNA structure by eIF4F. However, under hypoxic conditions, eIF4E2 may promote translation of specific mRNAs via bridging 5’-cap to the HIF-2α and RBM4 protein complex bound to the RNA hypoxia response element (rHRE) located in 3’-UTR of these mRNAs (41), (42). Hypoxia typically results in reduced protein synthesis rates by affecting the activity of the mTOR>4EBP signaling that regulates eIF4F complex formation for cap-dependent translation (43).

Currently, the role of SARS-CoV-2 Nsp2 protein in regulation of translation is not known. In this study, we demonstrated retention of Nsp2 on m^7^GTP-Sepharose presumably via interaction with eIF4E2 in cells grown under normal or hypoxic conditions. Moreover, we observed colocalization of Nsp2 with eIF4E2, 40S ribosomal protein S3 and with ER marker calnexin in human embryonic kidney HEK293T cells, suggesting the presence of Nsp2 in close proximity to the protein synthesis sites in ER. Finally, we demonstrated increased translation of capped and HCV-IRES-containing Luciferase mRNAs under normal and hypoxic conditions in cells expressing Nsp2. We propose that SARS-CoV-2 Nsp2 protein functions to reprogram translation machinery that would promote synthesis of viral and host proteins that may be beneficial to viral replication, assembly and/or secretion. It is interesting to speculate how the interaction of the Nsp2 protein with eIF4E2 may alter the translational machinery, and particularly during a patient’s condition of oxygen deficit, for its own program of viral protein synthesis.

## RESULTS

### Nsp2 and eIF4E2 co-purify on Biotin Agarose and m^7^GTP-Sepharose

To investigate the interaction between Nsp2 and eIF4E2 we first generated a cell line expressing SARS-CoV-2 Nsp2 protein containing an N-terminal Streptavidin tag. As a control cell line, we used HEK293T cells transfected with empty vector. We selected puromycin resistant Nsp2- and control cells in a similar manner. Western blot analysis confirmed expression of Strep-Nsp2 protein in the SARS-CoV-2 Nsp2 cells (**Fig. 1A**). Next, we evaluated interaction of Nsp2 and eIF4E2. Cell lysates containing equal amount of total protein from control and Nsp2-transfected cells were incubated with either Biotin-Agarose (**Fig. 1B**) or with m^7^GTP-Sepharose (**Fig. 1C**). With both affinity matrices, we could confirm the interaction between Nsp2 and eIF4E2 as the two proteins were enriched upon copurification only from extracts of cells expressing Nsp2.

**Figure 1.**
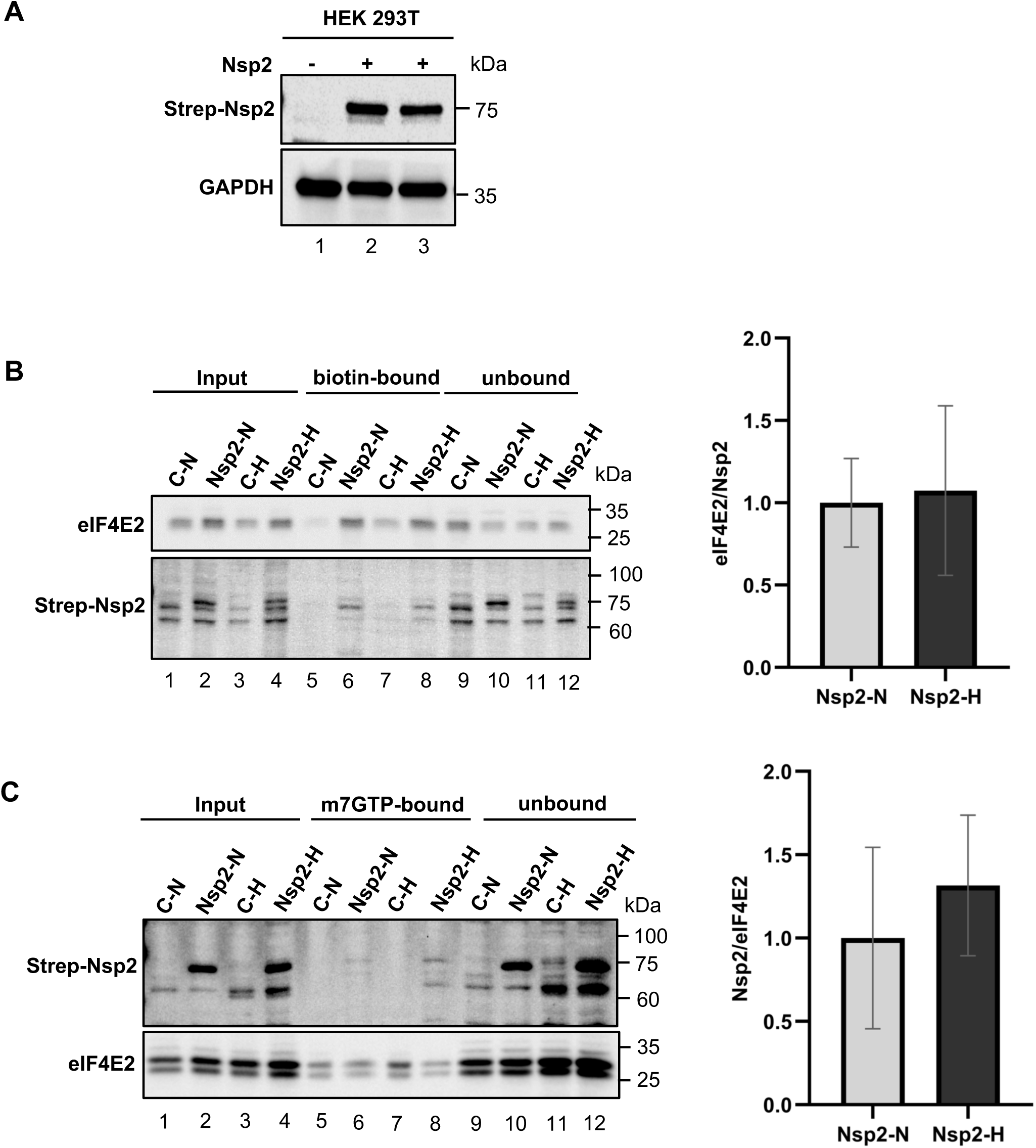
Expression of Nsp2 and its binding to eIF4E2. **(A)** Representative image of western blot analysis of expression of Strep-Nsp2 in HEK293T cells stably transfected with either Lipofectamine-3000 or LTX. (**B)** Biotin-Agarose pull-down. Left, western blot analyses of eIF4E2 (upper panel) and Nsp2 (lower panel). Right, graph represents densitometric ratio of eIF4E2 to Nsp2. (**C)** m^7^GTP-Sepharose pull-down. Left, western blot analyses of Nsp2 (upper panel) and eIF4E2 (lower). Right, graph represents densitometric ratio of Nsp2 to eIF4E2. C-control; N – normoxia; H – hypoxia. The blots shown are representative of three completely independent experiments.

### Cellular localization of Nsp2

Previous studies have shown that Nsp2 viral protein localizes in the cytoplasm of cells (44). To investigate whether Nsp2 co-localizes with eIF4E2 we performed immune-fluorescent staining of controls and Nsp2-expressing cells grown under normal or hypoxic conditions (**Fig. 2A**). We observed colocalization of Nsp2 and eIF4E2 in cell cytoplasm under both conditions. Next, we investigated whether Nsp2 present at the sites of protein synthesis. We observed colocalization of Nsp2 with the rough endoplasmic reticulum marker, calnexin (**Fig. 2B**) and with a small ribosomal subunit protein S3 (RbS3) (**Fig. 2C**). These data indicate that Nsp2 co-localizes with eIF4E2 at dense granular cytoplasmic structures that are also sites of active protein synthesis, and thus presumed to be RER. Interestingly, we observed increased co-localization of Nsp2 with eIF4E2, calnexin, and RbS3 under hypoxic conditions. It was reported that the synthesis of proteins targeted for export/secretion (RER-bound transcripts) are highly increased upon hypoxic conditions (45).

**Figure 2.**
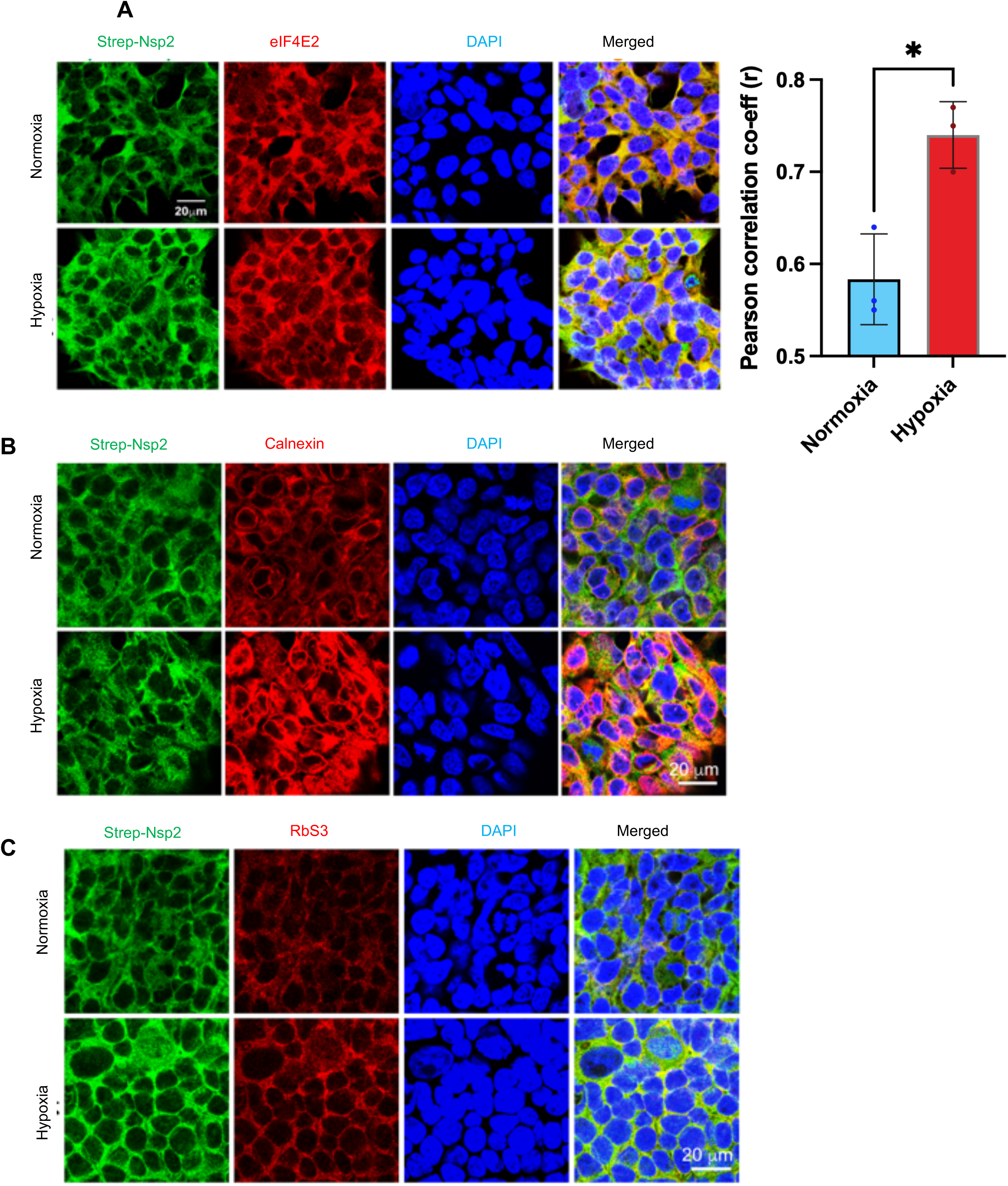
Nsp2 co-localizes with eIF4E2 and RER under normal and hypoxic conditions. **(A)** Strep-Nsp2 was probed using anti-strep antibody (green) and eIF4E2 probed using anti-eIF4E2 antibody (red), DAPI (blue) stained nuclei. Merge channel from max intensity projections of z-stacks under hypoxia shows increased co-localization (yellow). **(B)** Strep-Nsp2 (green) and calnexin (red) probed. Under hypoxia, granular staining in merged channel (yellow) shows increased co-localization from best focal plane. **(C)** Representative image of Ribosomal marker (RbS3) probed using anti-RbS3 antibody (red) and Strep-Nsp2 (green). Colocalization shown in merged channel. DAPI stained nuclei.

### Nsp2 stimulates cap- and HCV-IRES-dependent translation

To investigate whether Nsp2 affect protein synthesis, we monitored expression of Renilla and Firefly luciferases in cells grown under normal and hypoxic conditions (1% oxygen) for 12-18 hours. Control and Nsp2-cell were transiently transfected with a [pCMV Luc Renilla-HCV IRES-Firefly luciferase] plasmid that expresses Renilla in a cap-dependent manner and Firefly luciferase in an IRES-dependent manner (cap-Ren-HCV-FF). We observed that Nsp2 cells have higher expression levels of both Renilla and Firefly luciferases under normoxia and hypoxia (**Fig. 3**). To evaluate whether observed increase in translation in Nsp2-cells is due to more efficient transfection or due to stabilization of mRNA we performed real-time PCR to measure the levels of GAPDH, Renilla and Firefly mRNAs. We found that GAPDH mRNA levels were similar between control and Nsp2-cells in both normoxia and hypoxia. Ratio of GAPDH mRNA in control *vs* Nsp2-cells was 0.98 ± 0.02 (n=6) under normoxia and 0.99 ± 0.01 (n=6) under hypoxia. To analyze the level of Renilla and Firefly mRNA, we first normalized luciferase signal to the GAPDH in the corresponding sample. Then we have measured the ratio of luciferase mRNA (normalized to GAPDH) to the average of corresponding luciferase mRNA in control cells. We found that the expression level of Renilla mRNA was similar between control and Nsp2 cells in normoxia and hypoxia: 1 ± 0.13 in control cells (n=3) and 0.9 ± 0.32 in Nsp2-cells (n=3) under normoxia; and 1 ± 0.26 in control cells (n=3); and 1 ± 0.37 in Nsp2-cells (n=3) under hypoxia. We have also confirmed that the ratio of Renilla mRNA to Firefly mRNA is similar in control and Nsp2-cells cells in normoxia and hypoxia: 1.1 ± 0.11 in control cells (n=3) and 0.9 ± 0.12 in Nsp2 cells (n=3) under normoxia; and 1.2 ± 0.06 in control cells (n=3) and 0.99 ± 0.098 in Nsp2-cells (n=3) under hypoxia. These data suggest that Nsp2 expression does not affect transfection efficiency, level of expression, or stability of mRNAs driven from the [pCMV Luc Renilla-HCV IRES-Firefly luciferase] plasmid under normoxia or hypoxia.

**Figure 3.**
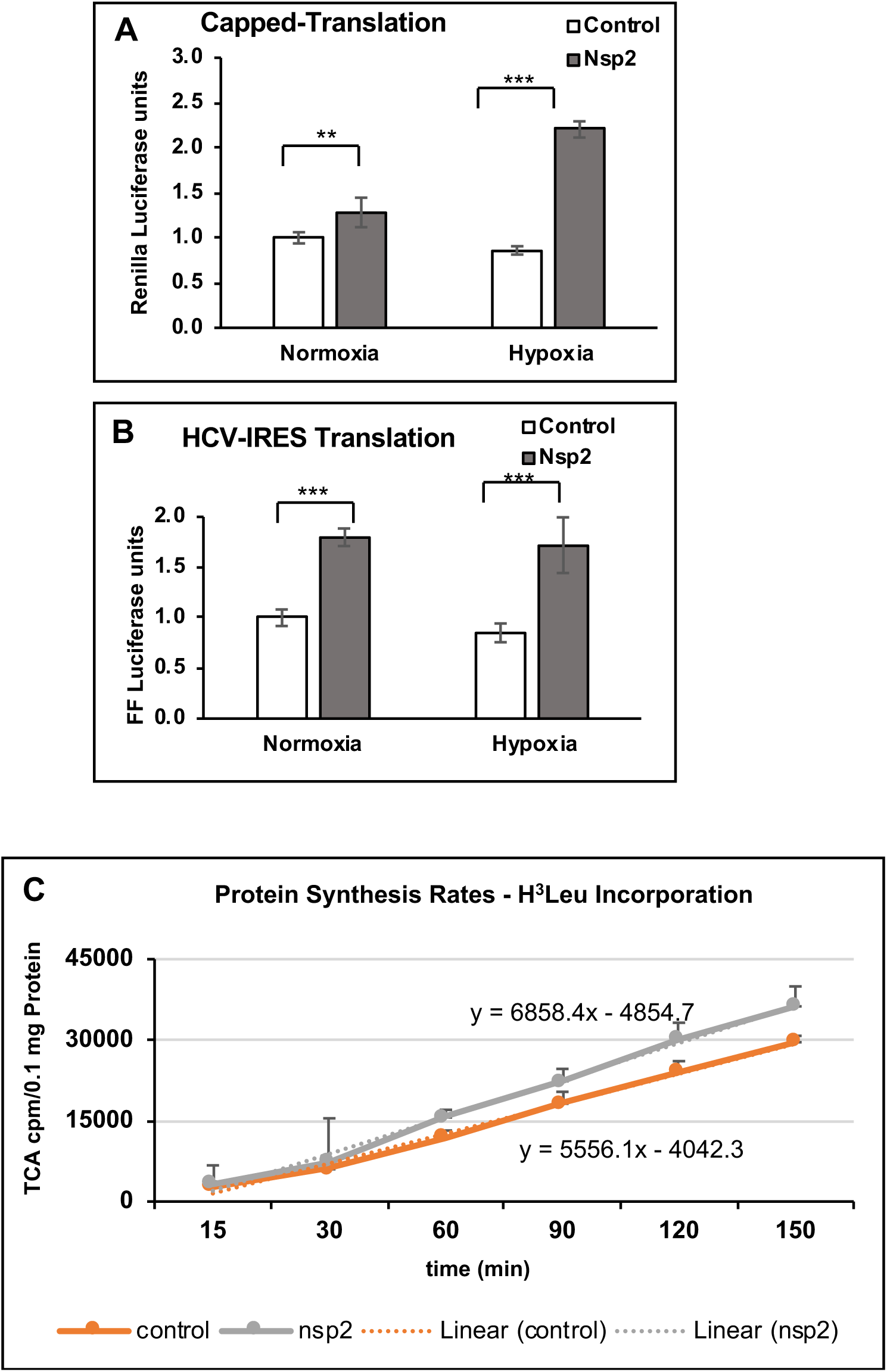
Nsp2-cells demonstrate higher cap- and HCV-dependent translation under normal and hypoxic conditions. Expression of Renilla and Firefly luciferases from capped-Renilla-HCV-IRES-FF Luciferase mRNA in control (white bars) and Nsp2 cells (grey bars) grown under normal or hypoxic conditions. Graphs represents mean of Renilla units (expressed from the capped mRNA), set as one **(A),** and Firefly (expressed from the HCV IRES mRNA) **(B)** luciferase signals (±SD, n=3). Statistical analysis of luciferase signals to one in control cells grown under normal or hypoxic conditions was performed using Student’s t test, with p < 0.05. **(C)** Nsp2-cells have higher translation rate. Graph represents the rate of H^3^-Leu incorporation in 0.1 mg of recovered protein extract. The difference in the slopes of linear protein synthesis between the two cell lines was highly significant (p = 0.0157 t-test).

### Nsp2 stimulates the rate of translation

To evaluate the rate of protein synthesis (PS) we monitored the rate of H^3^-Leu incorporation in the control and Nsp2-cell lines grown under normoxic conditions. We determined that there is statistically significant greater PS rate in the Nsp2-expressing cells compare to that in control cells (Fig. 3C). We did not measure the rate of PS under hypoxic conditions due to the technical difficulties to do such experiment in the hypoxic chamber because of the difficulty of adding the label after hypoxia is reached, and then for removing the time aliquots. Increased rate of PS in Nsp2-cells indicate that Nsp2 expression affects the rate of but does not distinguish between cap- and IRES-dependent translation.

### Nsp2 stimulates translation of eIF4E-dependent mRNAs

To investigate how expression of Nsp2 affects translation of endogenous mRNAs we performed polysomal analysis of cells grown under normal and hypoxic conditions. First, we observed increase in 80S peak in control cells grown under hypoxia compared to that in normoxia confirming inhibition of initiation step of translation (**Fig. 4A**). At the same time, under both conditions, Nsp2 cells had lower 80S ribosomal peak compared to that in control cells, suggesting more efficient initiation step of translation. To evaluate the effect of hypoxic conditions on the distribution of ribosomes along sucrose density gradients, we isolated RNA from each fraction of the sucrose density gradients and performed real-time PCR assay using specific primers for 18S ribosomal RNA. We then calculated a ratio between 18S RNA co-sedimented with polyribosomes (fractions #7-11) and monosome (fractions #4 and 5) (Fig. 4A, Ps/Ms). We observed that while in control cells Ps/Ms ratio decreased from 3.8 ± 0.2 under normoxia to 2.3 ± 0.3 under hypoxia, in Nsp2-cells the ratio Ps/Ms stayed higher under both conditions: 4.2 ± 0.4 and 4.3 ± 0.3 under normoxia and hypoxia, respectively. We also calculated a ratio between 18S RNA co-sedimented with heavy polyribosomes, containing efficiently translated mRNAs (fractions #7–11), and light polyribosomes, containing inefficiently translated mRNAs (fractions #1-6) (**Fig. 4A**, H/L). We observed that in control cells, the H/L number decreased from 0.7 ±0.1 observed under normal conditions to 0.5 ± 0.1 under hypoxia suggesting enrichment of 18S in light polysomal fractions, in agreement with inhibition of initiation step of translation. In contrast, in Nsp2 cells, the 18S H/L number did not differ between normoxia and hypoxia, 1.0 ± 0.2 and 1.1 ± 0.1, respectively. Higher Ps/Ms and H/L ratios in Nsp2-cells suggest a more efficient translation under both normal conditions and hypoxia compared to the control cells.

**Fig 4.**
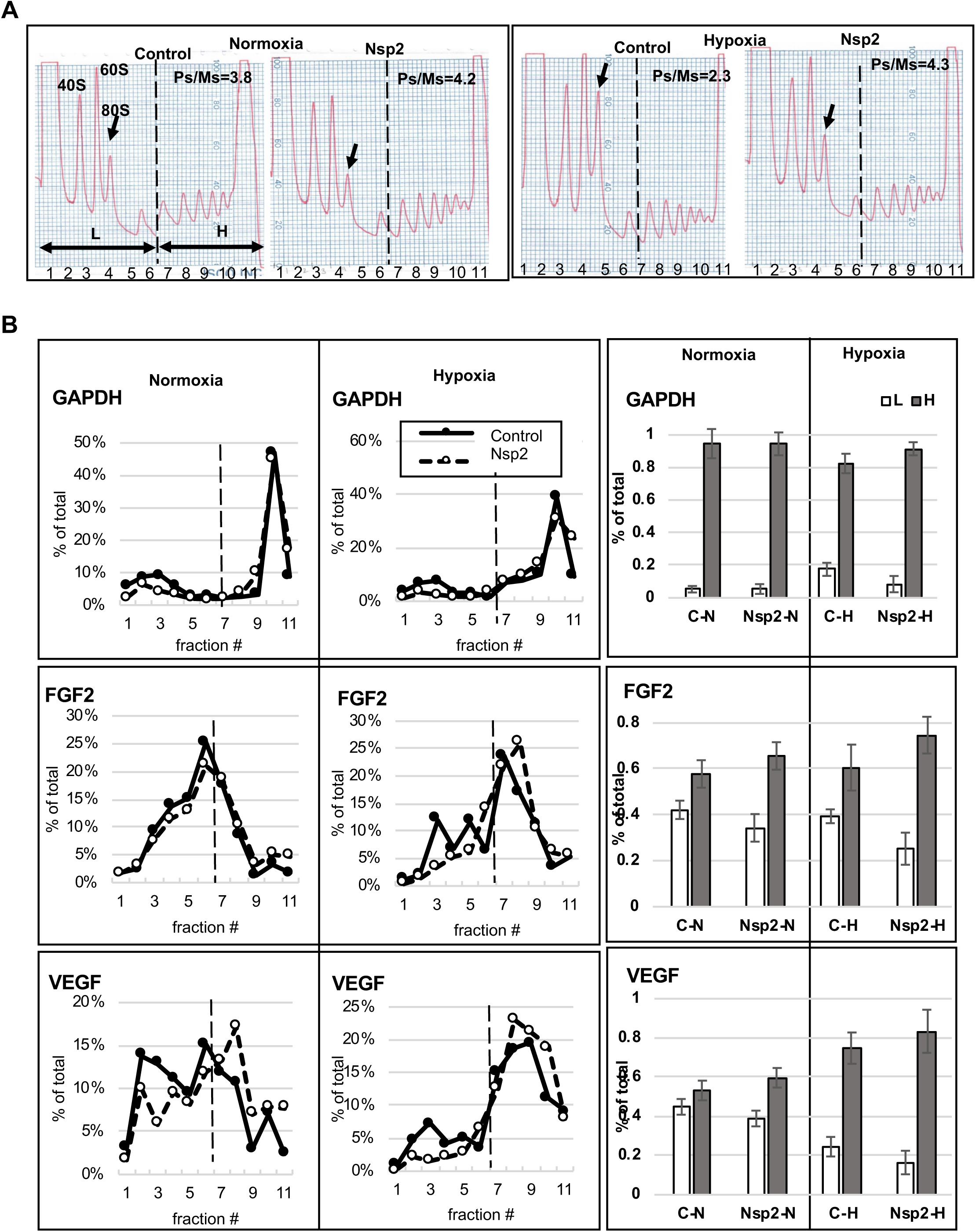
Polysomal analysis of control and Nsp2-cells in sucrose density gradients. **(A)** The representative of A254 profiles representing lysates distribution of cells grown under normal (left panel) and hypoxic (right panel) conditions in sucrose density gradients. Top of the gradient is on left side of each image. Position of 40S, 60S, and 80S along Ps profile are indicated. Arrow, 80S pick. L, light polysomes representing low translated mRNAs, fractions #1-6; H, heavy polysomes representing more efficiently translated mRNAs, fractions # 7-11. Ps/Ms, ratio of the 18S RNA signals in polysomes vs monosomes (fractions #4 and 5). **(B)** Real-time-PCR analysis of GAPDH, FGF2, and VEGF-C mRNAs distribution in sucrose density gradients. Left panels, distribution of the corresponding mRNA along sucrose density gradient similar to ones presented on (A). Graphs represent percentage of individual mRNA in each fraction. Filled circle, control cells; empty circle, Nsp2-cells. Right panels, graphs representing percentage of individual mRNA in light polysomal fractions #1-6 (white bars) and in heavy polysomal fractions #7-11 (grey bars) relative to the amount of this mRNA in all eleven fractions. The graphs represent the mean value from three independent polysomal ultracentrifugations (±SD).

To investigate the effect of Nsp2 expression on translation of the mRNAs that are specifically regulated at the initiation step of translation, we monitored the distribution of FGF2 and VEGF-C mRNAs in the sucrose density gradients. The FGF2 mRNA has a G-C rich 5’ highly structured region with a calculated stability of 50 kcal/mole and is known to be translationally stimulated by high level of eIF4E and eIF4F (46). Similarly, VEGF-C mRNA was shown to be translationally stimulated in the presence of high eIF4E level (47); (48). As a control, we also monitored distribution of GAPDH mRNA along sucrose density gradient, since it encodes a house-keeping protein and is not translationally regulated under hypoxia. First, we confirmed that neither Nsp2 expression nor hypoxia affect the total amount of GAPDH, FGF2, and VEGF-C mRNAs (data not shown). In **Fig. 4B**, graphs on the left, the amount of individual mRNA in each fraction is presented as a percentage of the amount of this mRNA in all 11 fractions. To analyze the amount of individual mRNA in the light (L, fractions #1-6) and heavy (H, fractions #7–11) polysomal fractions, we expressed it as a percentage of mRNA sedimented in all fractions set as 100 % (**Fig. 4B**, bars on the right). We did not observe any changes in GAPDH mRNA distribution along the sucrose density gradient in both, control and Nsp2 cells under normal and hypoxic conditions (**Fig. 4B**, upper panels). Statistical analysis showed that about 90% of the GAPDH mRNA co-sedimented with polyribosomal complexes (fractions #7–11) in all samples. FGF2 mRNA has similar polyribosomal distribution between control and Nsp2 cells under normal conditions (58±6% vs 66±6% in H fractions, respectively). However, under hypoxia, the percent of FGF2 mRNA in Nsp2 cells slightly increased in heavy polysomal fractions (75±8%) compared to that in control cells (61±10%) (**Fig. 4B**, middle panels) suggesting translational upregulation. Hypoxia increased the amount of VEGF-C mRNA in heavy polysomal fractions in both, control (75±6% in hypoxia *vs* 53±5% in normoxia) and Nsp2 (83±11% in hypoxia *vs* 60±5% in normoxia) cells (**Fig. 4B**, lower panels). These data suggest that VEGF-C mRNA is more efficiently translated under hypoxic conditions in control and Nsp2 cells. To confirm translational activation of FGF2 and VEGF-C mRNAs we monitored the FGF2 and VEGF-C protein expression and compared it to the GAPDH level (**Fig. 5**). The FGF2 mRNA has a G-C rich 5’ highly structured region containing at least two translation initiation sites. Translation from the cap-proximal CUG1 of the mammalian FGF2 mRNA was shown to be stimulated by excess eIF4E/eIF4F *in vitro* and *in vivo* (46, 49). In contrast, translation from the internal AUG (~400 nt downstream from it) is believed to occur via an IRES-driven mechanism (50), particularly under hypoxic conditions, although there is also some evidence that under presence of excess eIF4F, it may be translated via leaky-scanning upon melting of the highly G/C-rich secondary structure of the ~500 nt-long 5’UTR (46, 49). Although most evidence for this complex mechanism of translation initiation at alternative CUG and AUG start codons is attributed to features in the 5’UTR, one should not disregard the demonstrated potential regulation via the ~6000 nt long 3’UTR (49, 51). We found that in hypoxic conditions, the translation from CUG1 was increased in both control and Nsp2 expressing cells (**Fig. 5A, left image**). Under milder hypoxia, translation from the CUG start codon was highly stimulated in the Nsp2 but not in control cells (**Fig. 5A, right image** and quantified from independent experiments in **Fig. 5B**). In agreement with the VEGF-C mRNA shift to the heavy polysomes under hypoxia, we observed increased VEGF-C protein levels in control and Nsp2 cells under hypoxic conditions (**Fig. 5C and Supplemental Fig. 3**).

**Fig. 5.**
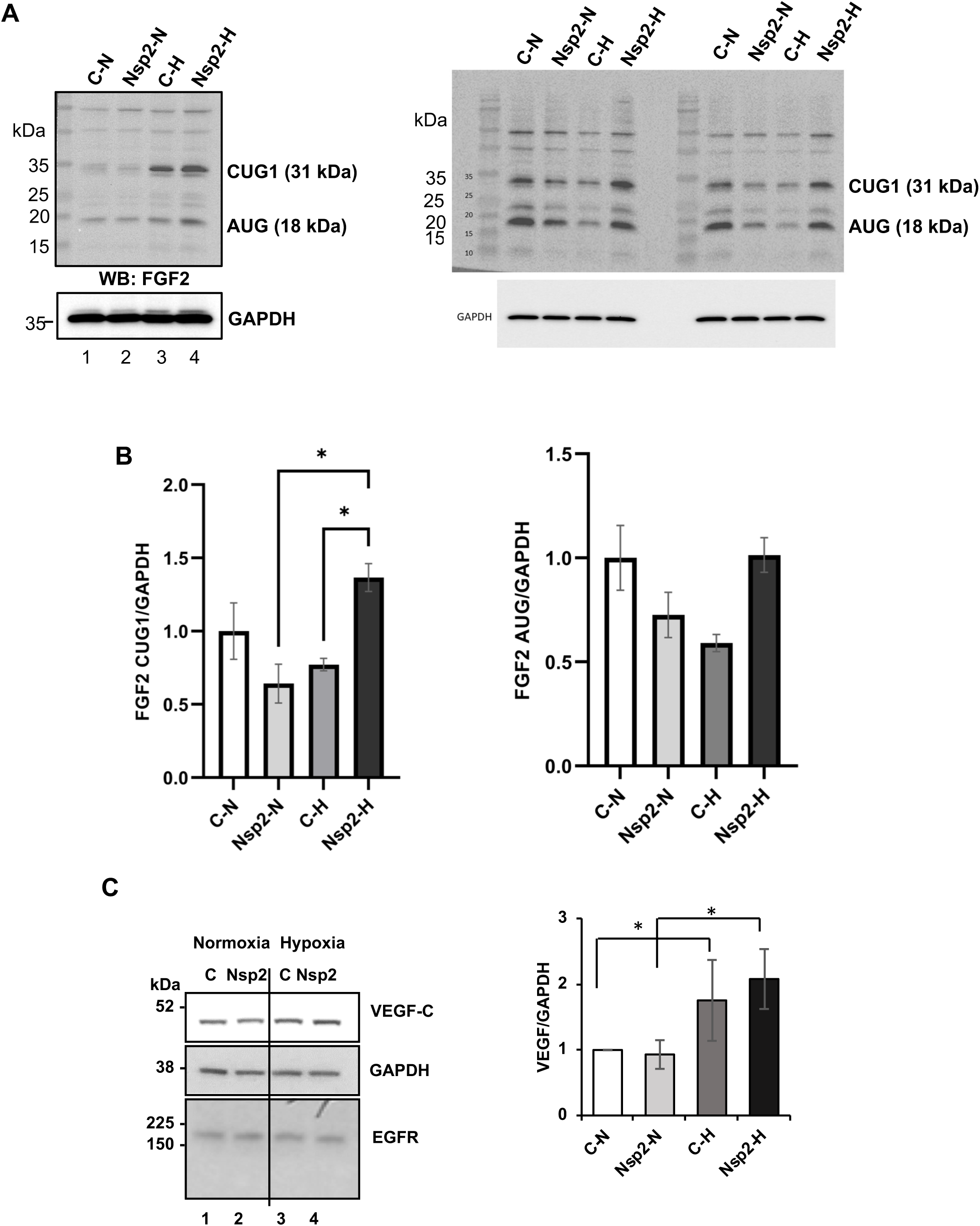
Western blot analyses of FGF2 and VEGF proteins in lysates. **(A)** FGF2 CUG1 and AUG isoforms in Nsp2 expressing cells in normoxic and hypoxic conditions (Right and left panel represents three biologically independent experiments). GAPDH was used as a normalization control **(B)** Densitometric ratio of CUG1 to GAPDH (left panel) and AUG to GAPDH (right panel). **(C)** Left, the representative images of western blots of VEGF-C, EGFR, and GAPDH. Right, graphs of the densitometric analysis of western blots. The graphs represent the mean ratio of signal of VEGF-C to GAPDH normalized to that in control cells grown under normal conditions, set as one (±SD). Western blot analysis was performed for each set of proteins using lysates from three separate experiments and repeated at least twice for each set of lysates.

### Nsp2 expression affects composition of translation initiation complexes

To investigate whether Nsp2 expression affects composition of translation initiation complexes we first analyzed proteins co-sedimented with 40S ribosomal peak in the sucrose density gradients. According to the polysomal profiles, fractions #2 contain 40S (**Fig. 4A**) but lack a biomarker of large 60S ribosomal subunit, RsL7a (**Fig. 6**). This suggest that proteins in fractions #2 may be a part of 48S initiation complexes. Western blot analysis revealed that under normal conditions, both control and Nsp2 cells contain eIF4G and eIF4A in their fractions #2 of polysomal gradient confirming the presence of initiation complex eIF4F (**Fig. 6A**). However, control cells also had eIF4E2 and 4EBP in fraction #2 suggesting the presence of some inhibitory complexes. Hypoxia increased the 4EBP signal in fractions #2 in both control and Nsp2 cells but at a lesser extent in the Nsp2 cells (**Fig. 6B**). These data suggest that Nsp2 expression reduces the inhibitory effects of eIF4E2 and/or 4EBP on translation. Indeed, Nsp2 cells exhibit stronger eIF4G signal along polysomal fractions in both normal and hypoxic conditions compared to control cells suggesting more efficient initiation of translation (**Fig. 6**, eIF4G). Interestingly, we detected Nsp2 signal in the fractions #2 under normal and at a lesser amount in hypoxic conditions. This result suggests that Nsp2 protein may be a part of complexes that regulate initiation step of translation.

**Fig. 6.**
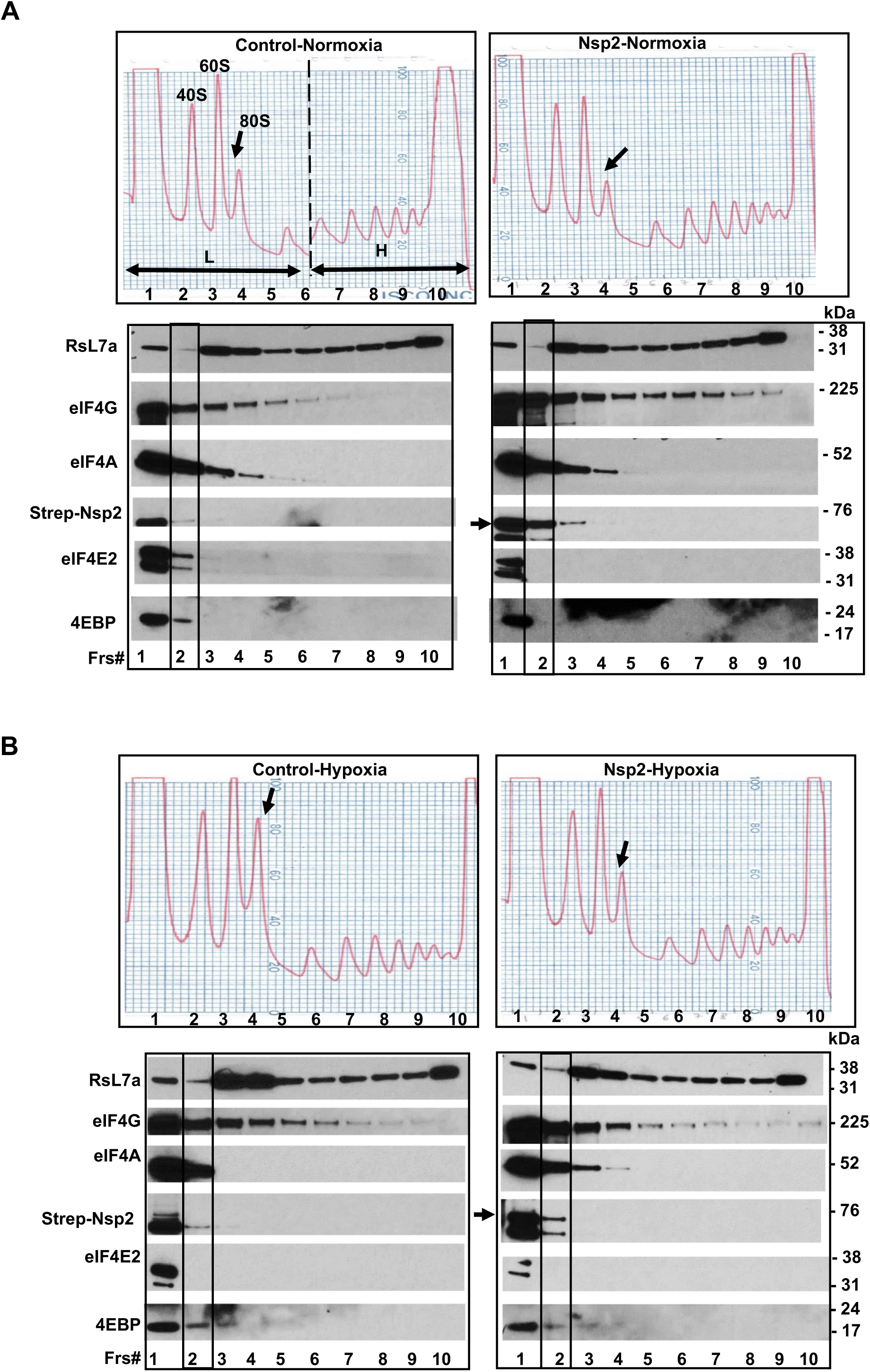
Western-blot analysis of polysomal fractions. The representative images of A254 profiles (as in Fig. 4A) and western blots of 60S ribosomal protein L7a (RsL7a), eIG4G, eIF4A, Strep-Nsp2, eIF4E2, and 4EBP in corresponding fractions (Frs) along sucrose gradients in control and Nsp2 cells grown under normal **(A)** and hypoxic **(B)** conditions. The western blot analysis of proteins was investigated in three sucrose density gradients using lysates from separate experiments. Arrow, position of Nsp2 signal.

### Alterations in eIF4F components in hypoxic conditions and in presence of Nsp2

First, we confirmed that our hypoxic conditions induce the biomarkers of hypoxia by probing lysates with anti-Hif-1α antibodies. Indeed, we observed increase in Hif-1αα expression in both control and Nsp2-cells under hypoxia (**Fig. 7A**). Hypoxia is also known to downregulate protein synthesis largely through changes in mTOR and eIF2α activities (43, 45). Indeed, mTOR phosphorylation at S2448 (a marker of its activity) was suppressed under hypoxia, in both controls and Nsp2-expressing cells (**Fig. 7A**). Consequently, there was a loss of 4EBP phosphorylation, both seen by P-S65 and increase of the high mobility form after hypoxic exposure. Although the ratio of P-4EBP to 4EBP was decreased in both control and Nsp2 cells, overall Nsp2 cells maintained a higher ratio of P-4EBP to 4EBP (**Fig. 7A, graph on a right**). This is expected to be reflected in the association of 4EBP with eIF4E, thereby releasing eIF4E for its recruitment in the eIF4F complex.

**Fig. 7.**
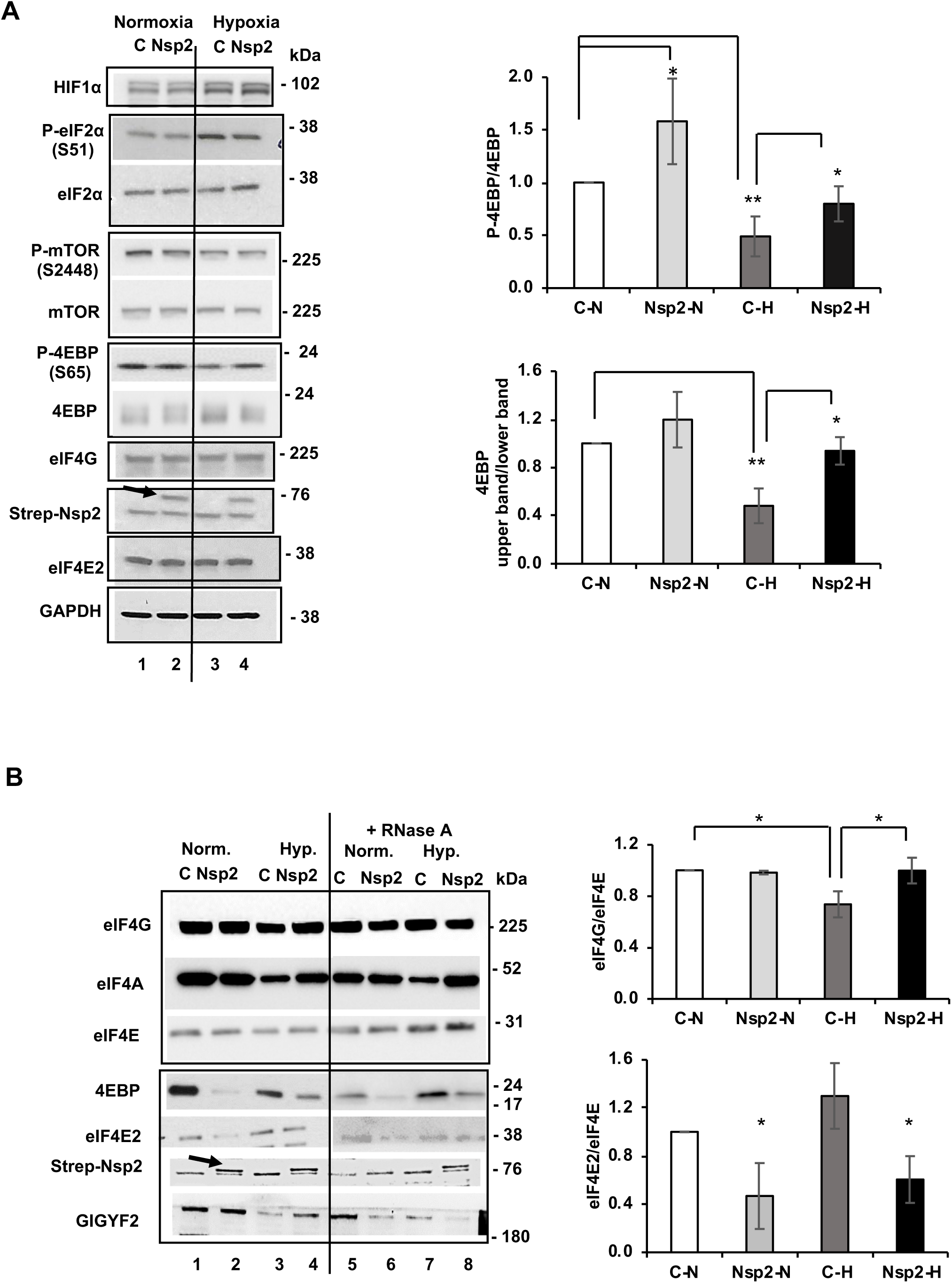
Nsp2 may affect composition of the cap-binding complexes. **(A)** Left, the representative images of western-blot analyses of proteins expressed in control and Nsp2-cells under normal and hypoxic conditions. Arrow indicates position of Strep-Nsp2. Right, ratio of densitometric signal of P-4EBP to total 4EBP in cells grown under normal or hypoxic conditions normalized to that in control cells grown under normal conditions, set as one (±SD, n=3). **(B) Left,** the representative images of western blot analysis of proteins bound to m^7^GTP-Sepharose under normal and hypoxic conditions in control and Nsp2-cells. Arrow indicates position of Strep-Nsp2. Right, ratios of densitometric signals of eIF4G (upper graph) and eIF4E2 (lower graph) to eIF4E bound to m^7^GTP-Sepharose in experiments presented in lanes 1-4 (±SD, n=3). Graph eIF4E/eIF4G: p = 0.014 for C-H *vs* C-N; p = 0.033 for Nsp2-H *vs* C-H. Graph eIF4E2/eIF4E: p = 0.03 for C-H *vs* C-N; p = 0.025 and 0.023 for Nsp2-H *vs* C-N and *vs* C-H, respectively.

To investigate the composition of proteins associated with the cap-binding complexes, we performed a pull-down experiment using m^7^GTP-Sepharose. To avoid contribution of RNA into the complexes retained on m^7^GTP-Sepharose we incubated some samples with RNase A. As we expected, we observed lower level of 4EBP associated with m^7^GTP-Sepharose (meaning decreased binding with eIF4E (1) that would correspond in its greater association with eIF4G as seen from their ratio) in Nsp2-cells compared to that in control cells under both, normoxic and hypoxic conditions regardless of RNase A treatment (**Fig. 7B**). This observation can explain in part the retention of active, cap-dependent, protein synthesis under hypoxia in cells expressing Nsp2. The alternative explanation involving the activity of eIF2α seems less likely since it was hyperphosphorylated equally in control and Nsp2 cells after hypoxia (**Fig. 7A**). Instead, we noticed that eIF4E2 had a lower retention on m^7^GTP-Sepharose when Nsp2 is expressed, suggesting that Nsp2 interferes with its cap-binding capacity (**Fig. 7B** and **Fig. 1C**). This is an intriguing observation, suggesting that even though Nsp2 would have no direct effect on the eIF4F complex formation and retention on m^7^GTP-Sepharose, under conditions where eIF4F and eIF4E2/RBM4 compete for binding to the pool of capped-mRNAs, the presence of Nsp2 could shift the overall equilibrium towards eIF4F binding to mRNAs and promote translation. We should add that we have no evidence confirming the Uniacke’s reports that eIF4E2 promotes translation of select mRNAs containing RBM4 binding sites under hypoxic conditions, leading for instance to increased EGFR expression (52); see Fig. 5c). Our evidence is rather consistent with the previous reports that eIF4E2 acts as a competitive inhibitor of eIF4E and thus translation. In fact, depletion of eIF4E2 with siRNA in Hela cells was reported to result in a 30% increase in protein synthesis rates (33), and we obtained similar results with the lung epithelial carcinoma cells A549 (Supplemental Figure 1) confirming its general inhibitory mechanism even when applied to the more relevant lung tissue in the context of SARS-CoV-2 NSP2-eIF4E2 interaction.

## DISCUSSION

In our study, we observed that Nsp2 protein interacts with eIF4E2 under normal and hypoxic conditions. Our confocal microscopy data indicates that eIF4E2, which may serve as an alternative translation initiation factor under stressed condition, co-localizes preferentially with Nsp2 (viral protein) under hypoxia. As localization of a protein implies its site of function, we asked if the Nsp2 protein harbors at the rough endoplasmic reticulum which is the cellular localization of translation machinery. We found that indeed, Nsp2 has prominent co-localization with calnexin (RER marker). Further to confirm the Nsp2 association at ribosomal compartments, we found that there is a tendency to conglomerate in ribosomal units (RbS3) upon hypoxia. A study analyzing localization of SARS-CoV-2 proteins in HEp-2 cells demonstrated that SARS-CoV-2 proteins localized either in cytoplasm (Nsp2, Nsp3C, Nsp4, Nsp8, Spike, M, N, ORF3a, ORF3b, ORF6, ORF7a, ORF7b, ORF8, ORF9b and ORF10) or both in cytoplasm and nucleus (Nsp1, Nsp3N, Nsp5, Nsp6, Nsp7, Nsp9, Nsp10, Nsp12, Nsp13, Nsp14, Nsp15, Nsp16, E and ORF9a). Some of these proteins demonstrated colocalization with Golgi apparatus (Nsp15, M, ORF6 and ORF7a) or with ER (Nsp6, ORF7b, ORF8 and ORF10) (44). In our study, we observed colocalization of SARS-CoV-2 and Nsp2 protein RER marker calnexin, suggesting involvement of Nsp2 in regulation of protein synthesis and preferentially with the secretome based on localization.

We demonstrated that Nsp2-expressing cells have more efficient initiation of protein synthesis, probably via regulation of the eIF4E/eIF4F level. The presence of 5’-cap on genomic and sub-genomic SARS-CoV-2 mRNAs indicates the requirement of eIF4F for initiation of translation. Indeed, it was demonstrated that translation of sub-genomic SARS-CoV-2 mRNAs is cap-dependent and was shown to be highly sensitive to the eIF4E level (53). However, translation of genomic SARS-CoV-2 RNA does not require eIF4E and occurs in a cap-independent manner. It was demonstrated that the inhibitor that prevents the interaction between eIF4E and eIF4G also inhibited replication of human coronavirus HCoV-229 (54) suggesting the importance of the cap-dependent mechanism of translation for the viral cycle. Other mechanisms of initiation of SARS-CoV-2 coding regions include upstream ORFs, internal in-frame as well as out-of-frame ORFs and leaky scanning (55).

Translation initiation is a mostly regulated step whereby specific sets of mRNAs are selected from the pool of transcripts and recruited to the translating polysomes. Polysomal analysis of mRNAs preferentially translated in the Nsp2-expressing compared to control cell revealed FGF2 and VEGF-C mRNAs. FGF2 represents a unique case for translation, as its long and structured 5’UTR normally acts as a IRES dictating initiation from what is considered the standard ~18 kDa, AUG-start codon ORF, encoding a protein that is not secreted. However, when the activity of eIF4E/eIF4F is elevated, the excess helicase can unwind the secondary structure at the 5’UTR and promote initiation from the cap-proximal CUG1 or CUG2, resulting in co-linear extensions of the ORF that are much better secreted and have greater mitogenic activity (46). Similarly, it was demonstrated that VEGF-C expression is translationally up-regulated by high level of eIF4E and thus eIF4F (47), (48). Together, FGF2 and VEGFs are considered among the most powerful mitogenic factors for endothelia and other cell types and are typically produced upon tissue damage resulting in local hypoxia (56). These data suggest that Nsp2 stimulates translation of eIF4F-dependent mRNAs. A recent study analyzing ribosomal occupancy of host and viral mRNAs after SARS-CoV-2 infection of Vero E6 and Calu3 cells revealed low translational efficiency of viral transcripts that was comparable to the weakly translated host mRNAs (55). Low translational efficiency of viral transcripts was reported in another study of SARS-CoV-2 translotome during early and late phases of viral infection of a human lung cell line (Calu-3) (57). It was suggested that even SARS-CoV-2 transcripts are not preferentially translated, the high level of viral RNA rather deplete available translational resources to compete with the host mRNAs (55). In our study, we observed lower level of 4EBP associated with m^7^GTP-Sepharose in the cells expressing Nsp2 suggesting a mechanism how capped-sub-genomic SARS-CoV-2 mRNAs may compete for the host translation initiation complexes. Interestingly, even hypoxia suppressed cap-dependent translation of Renilla mRNA in control cells, in agreement with observed increase in dephosphorylated 4EBP, in Nsp2 cells, cap-dependent translation increased under hypoxia compared to normoxia.

Recently, proteomic analysis identified another binding partner of Nsp2 along with a host translation initiation factor eIF4E2, GIGYF2 (2), (21), (22). It was demonstrated that the eIF4E2-GIGYF2 protein complex is involved in miRNA-mediated translational inhibition and degradation of tristetraprolin-targeted mRNAs (33), (58). It was demonstrated that direct binding of Nsp2 to GYGYF2 enhances the formation of 4EHP-GIGYF2 protein complex that inhibits translation of IFN1β mRNA (59). A 1-350aa region of Nsp2 has been identified to be required for binding to eIF4E2-GIGYF2 complex (60). Authors demonstrated that cells expressing Nsp2 had less inhibitory effect on expression of the reporter containing 3’-UTR of IFN1β mRNA compared to control cells (60). Thus, both studies suggest a role of SARS-CoV-2 Nsp2 protein in suppressing production of IFN-β suppressing the antiviral immune response. However, these studies offer two contradictory models of how Nsp2-eIF4E2 interaction affects translation initiation of capped mRNAs. According to the Xu *et al.* model, binding of Nsp2 to GIGYF2 enhances the interaction between GIGYF2 and eIF4E2, all bound to the cap-5’-end of mRNA (59). In contrast, Zou *et al.’s* model suggest that binding of Nsp2 to the eIF4E2-GIGYF2 protein complex prevents its binding to the 5’-end cap-structure (60). In our study, we observed increased translation of capped-Renilla luciferase mRNA in Nsp2 cells suggesting stimulation of cap-dependent translation. We also observed stimulation of HCV-IRES-driven translation of Firefly luciferase mRNA in cells expressing Nsp2. It is possible that the initiation of HCV-IRES occurred via a leaky scanning/re-initiation mechanism in our conditions, or a hybrid, partly cap-driven mechanism, called Internal Ribosome Repositioning that was first identified in c-Myc mRNA (61).

Our current favored model is that Nsp2 acts to sequester a fraction of eIF4E2, which under normal and hypoxic conditions, is believed to be a competitive inhibitor for the formation of eIF4F complex and its binding to the 5’-cap, resulting inhibition of translation (42). In other words, Nsp2 acts as an inhibitor of the competitive inhibitor (eIF4E2) of eIF4F, thereby clearly stimulating cap-dependent translation, but also likely IRES driven as some still benefit from some functions of eIF4F. The virus may have evolved such function particularly to maintain active protein synthesis despite the hypoxic conditions that can arise during severe infection resulting a loss of lung function, in order to maintain expression of viral and at least a subset of cellular proteins. Studies in infected Primary Bronchial epithelial cells by ribosomal profiling have shown that SARS-Cov2 impact on translation was subtle (not at all diminished) and that the viral proteins were generally not translated better than cellular proteins (62); but none of this work carried out in hypoxic conditions. The Nsp2 protein of SARS-CoV-2 was shown to be significantly divergent from that of SARS-CoV, and it is highly likely that it has acquired novel functions that in this context cannot be studied in less dangerous but similar coronaviruses. Considering that SARS-CoV-2 Nsp2 is under selective pressure, it was suggested that mutation in Nsp2 protein could account for SARS-CoV-2 high ability of contagious and plays a key-role in the viral pathogenicity (26).

Our basic understanding of the mechanism of the Nsp2-mediated exploitation of eIF4E2-translation initiation and of the prevarication of viral proteins will be critical in assessing the soundness of certain proposed therapies. For example, already in the study by Gordon et al (2), ribavirin was suggested as a potential drug. Ribavirin was reported to work as a cap mimetic (63) and has led to inhibition of eIF4E-mediated progression of AML (64), and recently has been touted as effective in treatment of Covid-19 in preliminary clinical trials (65, 66). Identification of Nsp2 role in translation could lead to development of therapeutic treatment that suppress viral mRNA translation or curtail overall protein synthesis output that ultimately leads to cellular collapse and release of more virus.

## ACKNOWLEDGMENTS

Research reported in this publication was supported by grants from the FWCC Covid19 IDEA grant to NK and ADB and DoD-PCRP grant W81XWH-17-1-0417 to ADB. We thank Dr. David Gordon for providing the pLVX-EF1α-IRES-Nsp2 plasmid, Dr. Sarnow’s lab for providing the pCMVLucR-HCVLucF Luciferase plasmid, Dr. Hans Trachsel, Bern, Switzerland for anti-eIF4A antibodies, Dr. First for assisting with the plasmid preparation, Brian Latimer and the INLET Core Facility for assisting with live cell analysis using Cellomics ArrayScan™, the Research Core Facility at LSU Health Shreveport for assisting with the real-time PCR, confocal microscopy, and Dr. Shen and the COBRE Redox Core for assisting with the hypoxic chamber.

## AUTHOR CONTRIBUTIONS

N.K. and A.D.B. conceived and designed the study. N.K., M.I K., I.G., and R.F. performed the experiments. N.K., M.I. K., and I.G. generated the figures. All authors contributed to the manuscript during editing and rewriting phases.

## METHODS

### Cell culture and transfection

Human embryonic kidney 293T (HEK293T) cells were cultured in Dulbecco’s modified eagle medium (DMEM) supplemented with 10% FBS and 1% antibiotic-antimycotic and maintained in humidified incubator at 37 °C with 5% CO_2_. To generate Nsp2-cell line, HEK293T cells were transfected with pLVX-EF1α-IRES-Nsp2 mammalian construct (a kind gift from Dr. David Gordon, University of California, San Francisco) by Lipofectamine LTX and Plus^TM^ reagent following the manufacturer’s protocol. To generate control cells line, HEK293T cells were transfected with pDRGFP plasmid in which GFP is translated out of frame and does not produce a functional protein. Both control and Nsp2-cells were selected by 1 µg/ml of puromycin treatment for at least seven days. After the selection, cells were passaged, and early passages of cells were stored at −80°C. Cells underwent to selection by 0.1 µg/ml of puromycin after thawing for one more time and then grown without puromycin prior to the experiments.

### Fluorescence microscopy and image analysis

For immunofluorescence microscopy, sterile coverslips were placed into the wells of a 6-well plate and 0.3X 10^6 Nsp2 cells/well seeded 24hrs prior to hypoxia (1%O_2_) or normoxia treatment in 5%CO_2_ incubator at 37°C. Cells treated for 12hrs, and media removed. PBS wash twice. Cells fixed afterwards with 4%PFA in PBS at room temperature (RT) for 10mins. PBS wash done twice. Permeabilization was done using 0.5% Triton in PBS at room temperature for 10mins. PBS wash done twice. Blocking was done using PBS+1%BSA+0.1%Tween20 for 1hr at RT. Primary antibody (anti-strep, 1:500; anti-calnexin, 1:500; anti-eIF4E2, 1:100; anti-RbS3, 1:100) diluted in blocking buffer and added to samples. Incubated for overnight at 4°C. After one wash with PBS, secondary antibody (anti-rabbit and anti-mouse) against primary antibody was used for 1hr at RT. DAPI in anti-fade mounting media added on pre-cleaned slides and coverslips mounted. Fluorescence was detected and imaged using Nikon A1R-Super Resolution microscope equipped with a 60X oil objective lens (Apo 60x/1.40NA) with a numerical aperture of 1.4, two GaAsP detectors for laser 488nm and 561nm and a standard detector for 405nm (DAPI). Immunofluorescence images were taken using Nikon NIS-Elements C software at room temperature with 55 optical sections (n=3) separated by 0.1μm step-size. For representation of eIF4E2 and Nsp2 (main targets) colocalization was obtained for max intensity projection of 10 optical sections. For Calnexin and RbS3, best focal plane has been shown for representation.

### m^7^GTP-Sepharose and Biotin-Agarose pull-down assay

Control and Nsp2-expressing HEK293T cells were lysed in 0.3 ml of lysis buffer containing 50 mM Tris-Cl (pH 7.5), 100 mM KCl, 0.5% NP-), 2mM DTT, and EDTA free Halt™ Protease and Phosphatase Inhibitor Cocktail (100X – ThermoFisher Scientific). After sonication, cells were incubated on ice for 10 minutes and centrifuged at 14000 ×g for 10 mins at 4°C followed by the collection of supernatants and measurement of protein concentration by BCA Protein Assay Kit ThermoFisher Scientific. 200 µg of lysate was used for both m^7^GTP-Sepharose and Biotin-Agarose (Pharamcia Biotech) pull down experiments, which were diluted with equal volume of dilution buffer containing 50 mM Tris-Cl (pH 7.5), 2 mM DTT, and 5% glycerol. 200 µl of m^7^GTP-Sepharose and Biotin-Agarose (i.e., 50% slurry) were equilibrated in one ml of wash buffer containing 50 mM Tris-Cl (pH 7.5), 100 mM KCl, 2 mM DTT, and 5% glycerol for three times. Afterwards, all wash buffer was removed, and equal volume of dilution buffer was added to the resin. The diluted lysates from each experimental group were mixed with 50 µl of resin and incubated for 2 hours at 4°C with rotation. After the incubation, the resin was spun down and the supernatant was removed and saved as unbound fraction. After washing three times with 150 µl of wash buffer, proteins were eluted by resuspending the resins in 40 µl of 2X Laemmli (for m^7^GTP-Sepharose) or 4X Laemmli sample buffer (for Biotin-Agarose). The samples were heated at boiling temperature for 6 mins and further analyzed by western blotting.

### Protein Binding Assays on m^7^GTP-Sepharose (Fig. 7)

One hundred μg of lysates were prepared as described in the “Ribosome Fractionation” section and diluted in equal volume of buffer containing 50 mM Tris-HCl (pH 7.5) and 2 mM DTT. Then samples were mixed with 50 μl m^7^GTP-Sepharose, 50% slurry in buffer containing 20 mM Tris-HCl (pH 7.5), 100 mM KCl, 1 mM DTT, and 10% (v/v) glycerol. The resins were incubated for 2 h at 4°C with rotation. Duplicate samples were treated with RNase A for the last 40 minutes of incubation. Then, the resin was washed three times with 150-μl aliquots of the same buffer. Proteins were eluted in 20 μl 2x SDS-electrophoresis buffer and analyzed by Western blotting as it is described above. This experiment was repeated three times.

### Western blotting

For Fig. 1, samples from the pull-down experiments or cell lysates were run in either 7.5% or 12% SDS-PAGE gel and transferred to a PVDF) membrane. After blocking in 5% nonfat dry milk, the membranes were incubated in primary antibodies overnight at 4°C followed by the incubation in HRP-conjugated secondary antibodies for 1 h at room temperature. The blots were developed using ECL substrates and imaged in ChemiDoc Imaging System (Bio-Rad). Densitometric quantification of the blots were done using ImageJ software. For Figs. 5C and 7, to analyze the expression of proteins in cell lysates, equal amounts of total protein were loaded on a 12 % or 4–12 % NuPAGE® Novex® Bis–Tris Gel. The FullRange Rainbow protein molecular weight marker was loaded on the same gel to identify the position of specific proteins. Proteins were separated by SDS-PAGE gel and then transferred to a Nitrocellulose membrane using a Mini Trans-Blot cell. Expression of specific proteins in total lysates were determined by probing the membrane with antibodies against P-mTOR (Ser2448) (dilution 1:5000), mTOR (dilution 1:5000), P-4EBP (Ser65) (dilution 1:5000), P-eIF2α (Ser51) (dilution 1:2000), and eIF2α (D7D3) XP (dilution 1:4000), all from Cell Signaling; 4EBP (dilution 1:4000), eIF4E2 (dilution 1:1000), and Strep-Nsp2 (dilution 1:2000) all from Invitrogen; actin (dilution 1:4000) from Sigma; VEGF-C (dilution 1:1000) from R & D systems; and GAPDH (dilution 1:5000) from Fitzgerald. The membranes were incubated with primary antibodies in 5 % BSA in buffer TBS-T (20 mM Tris–HCl, 150 mM NaCl, and 0.1 % Tween 20, pH 7.5) overnight at 4 °C, washed three times for 15 min with TBS-T, and incubated for 1 h at room temperature with anti-mouse secondary antibodies or anti-rabbit secondary antibodies conjugated with horse-peroxidase in 5 % non-fat dry milk in TBS-T. Blots were developed with the Western Lightning ECL Pro development kit and exposed to HyBlot CL autoradiography film. Quantitative analysis of Western blot images was performed using the ImageJ software. Results are presented as the mean of two independent treatments using different cell passages. Student’s t test was applied to the data to determine statistical significance, and data with p value lower than 0.05 was considered to be statistically different.

### Translation of capped-Renilla and HCV-Firefly mRNA *in vivo*

The control and Nsp2 cells were cultured in DMEM supplemented with 10% fetal bovine serum and 1% penicillin-streptomycin. Cells were plated in to a 96-well white plate at 1×10^4^ cells per well and allowed to adhere overnight. Next day, cells were transfected with 0.1 µg DNA, pCMV Luc Renilla-HCV IRES-Firefly Luciferase, Lipofectamine LTX and Plus^TM^ reagent following the manufacturer’s protocol. After 9 hours of transfection, cells were washed, supplemented with fresh media and grown for another 12-18 hours either under normoxia or hypoxia (1% O_2_, 5% CO_2_, 37°C). Expression of Renilla and Firefly luciferases was detected using Dual-Glo® kit Luciferase Assay system according to the manufacture protocol. All Renilla luciferase signals (from capped mRNA) have been normalized to the mean value of Renilla in the control cells grown under normal conditions, set as 1. The experiment was repeated two times. All Firefly signals (from HCV-IRES) have been normalized to the mean value of Firefly luciferase in the control cells grown under normal conditions, set as 1. Student’s t test was applied to the data to determine statistical significance, and data with p value lower than 0.05 was considered to be statistically different.

### Evaluation of the rate of protein synthesis *in vivo*

10^5^ cells (control and Nsp2) were each plated in 12 wells of a 24-well plate with 1ml complete D-MEM and 10% FCS (duplicate samples). The next day the medium was replaced with medium containing 5µCi/ml L-[3,4,5-3H(N)]-Leucine (150 Ci/mmol - NEN), and sequential aliquots were removed at 1/2h intervals starting 15 minute later. After solubilization with 0.5 ml 1%SDS, 10% TCA insoluble material (0.1 mg Protein) was collected on 2.5 cm GFA filter. Following washing with 90% EtOH, the filters were air-dried and placed in scintillation vials for counting with OptiScint LLT NPE-Free Scintillation cocktail in a Beckman LS6500 counter.

### Ribosome Fractionation

The control and Nsp2 cells were cultured in DMEM supplemented with 10% fetal bovine serum and 1% penicillin-streptomycin. Cells were plated in to a 10 cm plates and allowed to adhere overnight. Next day, cells were either continued to grow under normal conditions or transferred to a hypoxic chamber (1% O_2_, 5% CO_2_, 37°C) for another 12-18 hours. Prior harvesting, cells were incubated with 50 μg/ml Cycloheximide for 10 minutes and then washed two times with warm PBS and one time with warm Ps wash buffer [50 mM Tris-HCl, pH 8.0; 140 mM NaCl; 5 mM KCl, 6 mM MgCl_2_], both buffers supplemented 50 μg/ml Cycloheximide. Cells were lysed with 0.3 ml per plate ice cold Lysis buffer [100 mM Tris-HCl, pH 8.0; 150 mM KCl, 10 mM MgCl_2_, 200 mM Sucrose; 5 mM DTT, 0.1 mg/ml Cycloheximide, 0.5 mg/ml Heparin, 0.5% Triton x-100, 0.5% NP-40, 0.5% Deoxycholate, 1x protease inhibitor EDTA-free, 1x phosphatase inhibitor 2, and 1x phosphatase inhibitor 3]. Lysates were centrifugated at 10,000 x g for 10 minutes at 4°C. Supernatants were aliquoted for polysomal and western blot analyses and stored at −80 °C until further use. Lysates of control and Nsp2 cells grown under normal or hypoxic conditions were analyzed by the polysomal ultracentrifugation. An equal amount of A_254_ optical units was layered onto a 15–45 % (w/v) sucrose gradient containing 50 μg/ml cycloheximide and 1 mM DTT and centrifuged in a Beckman SW41Ti rotor at 38,000 rpm at 4 °C for 2 h. Gradients were collected in 1-ml fractions with continuous monitoring of absorbance at 254 nm using an Isco syringe pump with UV-6 detector (Teledyne Isco Inc.). Samples were stored at −80 °C until further use. The polysomal analysis was repeated three times using lysates from different cellular passages. To analyze proteins along sucrose density gradients the 10 μl aliquots from each fraction were loaded on 4–12 % NuPAGE® Novex® Bis–Tris Gel, and analyzed by the Western Blotting as it is described above.

### RNA Isolation and Real-Time PCR

Before RNA isolation, 300 μl aliquots from each fraction after polysomal ultracentrifugation in sucrose density gradients were spiked with 100 pg of Luciferase mRNA (internal control, Promega). Then, RNA was purified with TRIzol®-LS reagent according to the manufacturer’s protocol. The RNA was further precipitated with 0.8 M Na-acetate and 1.2 M NaCl, re-suspended in RNase-free water and precipitated again with 2 M LiCl overnight at −20°C. Amplification and detection were performed using the iCycler IQ Real-time PCR detection system with Luna® universal One-Step RT-qPCR kit. Quantitative real-time PCR was used to measure the Firefly luciferase, 18S, GAPDH, FGF2, and VEGF-C RNAs level in each fraction. The 18S, GAPDH, FGF2, and VEGF-C RNAs levels were normalized with the luciferase internal control. Relative amount of individual RNA in each fraction (after normalization to Luciferase signal) was expressed as a percentage of the sum of this RNA in all 11 fractions set as 100 %. To assist statistical significance of the changes in the RNA redistribution along the sucrose density gradients, the percentage of individual RNA co-sedimented with light polyribosomes, containing inefficiently translated mRNAs (fractions #1-6) and heavy polyribosomes, containing efficiently translated mRNAs (fractions #7–11), was calculated as a percentage of the total mRNA. The percentage of individual mRNA in polysomal fractions was investigated in three polysomal analyses using lysates from different cellular passages.

### Polysomal analysis of proteins in sucrose gradient fractions

Expression of specific proteins in polysomal profiles (**Fig. 6**) was determined by probing the membrane with antibodies against RsL7a (dilution 1:4000) from Cell Signaling; 4EBP (dilution 1:4000), eIF4E2 (dilution 1:1000), and Strep-Nsp2 (dilution 1:2000) all from Invitrogen; mouse monoclonal anti-eIF4A antibody (dilution 1:8000) was a gift from Dr. Hans Trachsel, Bern, Switzerland; and rabbit anti-eIF4G antibody (dilution 1:20.000) was a gift from Dr. Robert E. Rhoads, LSUHSC. The 10 μl aliquots from each fraction after polysomal ultracentrifugation in sucrose density gradients were loaded on 4–12 % NuPAGE® Novex® Bis–Tris Gel, and analyzed by the Western Blotting as it is described above.

### Statistical analysis

Statistical analyses were performed using GRAPH-PAD PRISM 9 and MICROSOFT EXCEL software. Data are presented as mean± standard error of the mean (SEM). Statistical significance was determined by 2-tailed Student’s t-test when comparing the mean between two groups, or by one-way ANOVA followed by Tukey’s post hoc analysis when comparing more than two groups. P-values < 0.05 were considered significant.

## Key Resources Table

| REAGENT or RESOURCE | SOURCE | IDENTIFIER |
| --- | --- | --- |
| **Antibodies** |  |  |
| rabbit monoclonal anti-4EHP (eIF4E2) | Cell signaling Technology | Cat# 6916, RRID: AB_10839268 |
| mouse monoclonal anti-FGF2 | Santa Cruz Biotechnology | Cat# 145 |
| rabbit monoclonal anti-GAPDH | Cell signaling Technology | Cat# 2118, RRID: AB_561053 |
| mouse anti-strep | Invitrogen | Cat# MA5-17283 |
| rabbit anti-eIF4E2 | Invitrogen | Cat# PA5-110849 |
| rabbit Calnexin | Invitrogen | Cat# PA5-34754 |
| rabbit RbS3 | Cell signal tech | Cat# 2579S |
| goat anti-Mouse Alexa Fluor™ 488 | Life Technologies | Cat# A11029 |
| goat anti-Rabbit Alexa Fluor™ 594 | Life Technologies | Cat# A11012 |
| rabbit polyclonal anti-P-mTOR (S2448) | Cell signaling Technology | Cat# 2971,RRID: AB_330970 |
| rabbit monoclonal anti-mTOR | Cell signaling Technology | Cat# 2983,RRID: AB_2105622 |
| rabbit polyclonal anti-P-4EBP(S65) | Cell signaling Technology | Cat# 9451,RRID: |
|  |  | AB_330947 |
| rabbit polyclonal anti-4EBP (PHAS-1) | Invitrogen | Cat# 51-2900 |
| rabbit monoclonal anti-P-eIF2α (Ser51) | Cell signaling Technology | Cat#9721,RRID: AB_330951 |
| rabbit monoclonal anti-eIF2α(D7D3) XP | Cell signaling Technology | Cat#9721,RRID: AB_10692650 |
| mouse monoclonal anti-eIF4E2 (Figs. 6, 7, and S1) | Invitrogen | Cat#MA5-25412 |
| mouse monoclonal anti-GAPDH (Figs. 5C and S1) | Fitzgerald | Cat# 10R-G109A, RRID: AB_1285808 |
| goat polyclonal anti-VEGF-C | R & D systems | Cat# AF752-SP |
| rabbit polyclonal anti-Ribosomal protein L 7a (E109) | Cell signaling Technology | Cat# 2415,RRID:AB_2182059 |
| rabbit polyclonal anti-eIF4G | Gift from Dr. Rhoads (LSUHSC, Shreveport) |  |
| mouse monoclonal anti-eIF4A |  |  |
| rabbit polyclonal anti-Actin | Sigma | Cat# A2066, RRID: AB_476693 |
| Chemicals |  |  |
| Dulbecco's modified eagle medium (DMEM) | Sigma-Aldrich | D6429 |
| Antibiotic-Antimycotic (100X) | ThermoFisher Scientific | 15240-062 |
| Lipofectamine LTX and Plus<sup>TM</sup> reagent | ThermoFisher Scientific | 15338-100 |
| Puromycin | Sigma-Aldrich | P8833 |
| Non-idet P40 | Applichem | A2239 |
| Dithiothreitol (DTT) | Gold Biotechnology | 27565-41-9 |
| EDTA free Halt<sup>TM</sup> Protease and | ThermoFisher Scientific | 1861281 |
| Phosphatase Inhibitor Cocktail (100X) |  |  |
| m<sup>7</sup>GTP Sepharose | Pharmacia biotech | 27-5025-01 |
| Biotin Agarose | Agarose Bead Technologies | 4BCL-BI-5 |
| Laemmli Sample buffer (4X) | Bio-rad | 1610747 |
| PVDF Immobilon-P 0.45 mm | Milipore | IPVH00010 |
| Pierce™ ECL Western Blotting Substrate | ThermoFisher Scientific | 32106 |
| PAGE 20x running buffer | Invitrogen | Cat# NP0001 |
| PAGE 4-12% | Invitrogen | Cat# NP0322BOX |
| PAGE 10% | Invitrogen | Cat# NP0302BOX |
| Antioxidant | Invitrogen | Cat# NP0005 |
| Reducing agent | Invitrogen | Cat# NP0004 |
| Penicillin Streptomycin | Sigma | Cat# P4333-100ML |
| Nitrocellulose | We use Biotrace NT |  |
| Cycloheximide | VWR | Cat# 80059-088 |
| Phosphatase inhibitor 2 | Sigma | P5726 |
| Phosphatase inhibitor 3 | Sigma | P0044 |
| Heparin | VWR | 916-0 |
| Triton x-100 | VWR | EM9400 |
| NP40 | US Biological | N3500 |
| RNase-free H<sub>2</sub>O | Invitrogen | AM9939 |
| BPS | Boston BioProducts | BM-2205 |
| Sucrose | VWR | 97061-428 |
| Protease inhibitor | Roche | 12245300 |
| Centrifuge tubes for Polysomal analysis | Beckman | 331372 |
| Chloroform | VWR | IC0219400280 |
| Fetal Bovine Serum | Sigma | F4135-500ML |
| M<sup>7</sup>GTP-Sepharose (Fig. 7) |  |  |
| TRIzol™ LS Reagent | ThermoFisher Scientific | 10296028 |
| **Critical commercial assays** |  |  |
| BCA Protein Assay Kit | ThermoFisher Scientific | 23225 |
| Luna® universal One-Step RT-qPCR kit | New England BioLabs, Inc. | 103307-248 |
| Dual-Glo(R) Luciferase Assay System, E3005S | Promega | E2920 |
| **Software and algorithms** |  |  |
| ImageJ Software |  | RRID: SCR_003070 |
| GraphPad Prism | GraphPad | RRID:SCR_002798 |
| **Experimental models: Cell lines** |  |  |
| Hek293T cells | ATCC | CRL-1573 |
| **Recombinant DNA** |  |  |
| pLVX-EF1α-IRES-Nsp2 | A gift from Dr. David Gordon, University of California, San Francisco | N/A |
| pDRGFP | Addgene | 26475,<br>RRID:Addgene_26475 |
| **Other** |  |  |
| ChemiDoc Imaging System | Bio-rad | 12003154 |
| Nikon A1R Confocal microscope | Nikon | N/A |

## SUPPLEMENTAL INFORMATION

**Table S1.**
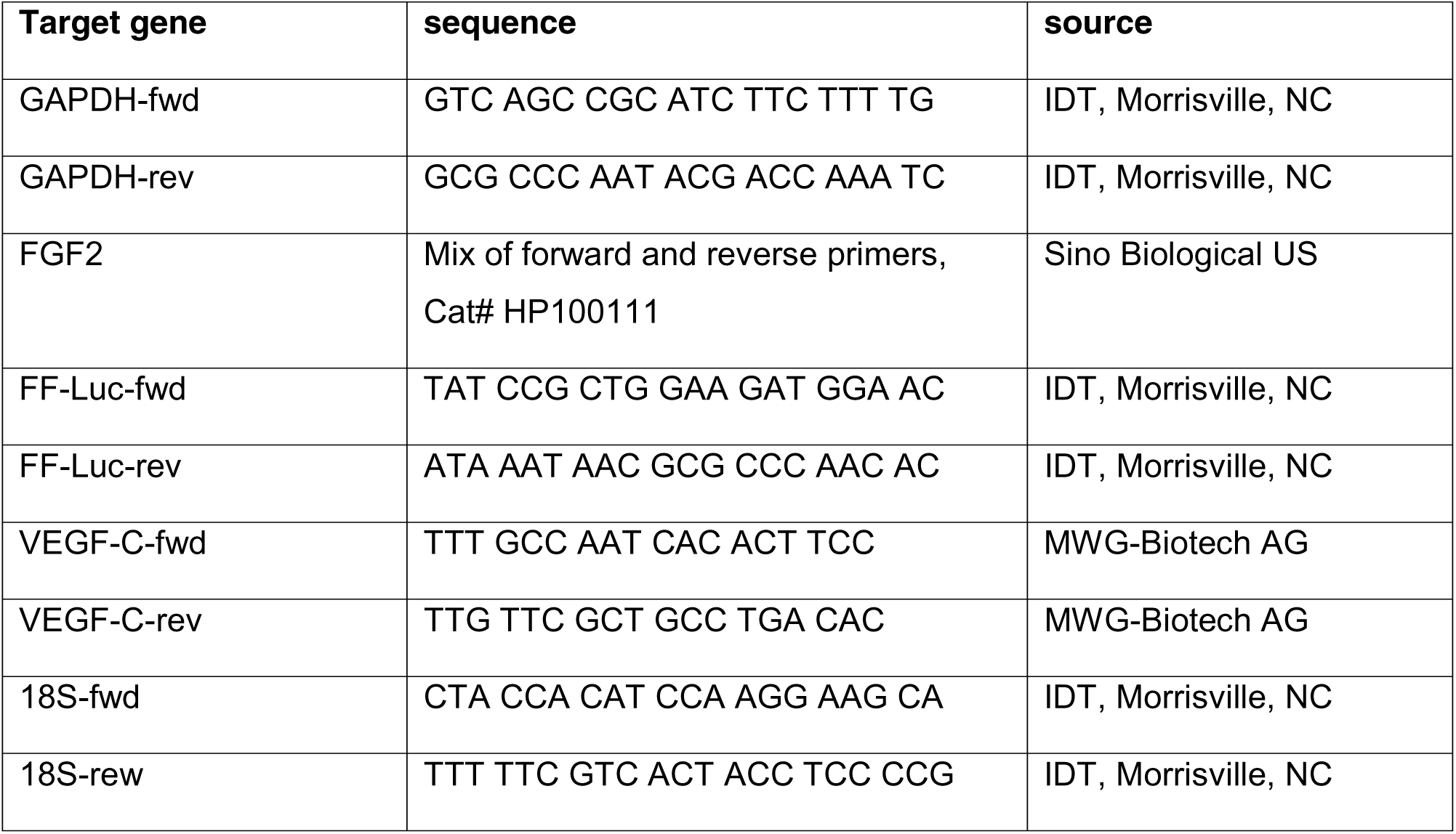
Primers used for real-time qPCR to analyze total RNAs and RNAs in polysomal gradients. Related to the STAR Methods.

