## Supplementary material for "SARS-CoV-2 Protein Nsp2 Stimulates Translation Under Normal and Hypoxic Conditions": SI Fig1-3

### Supplemental Information

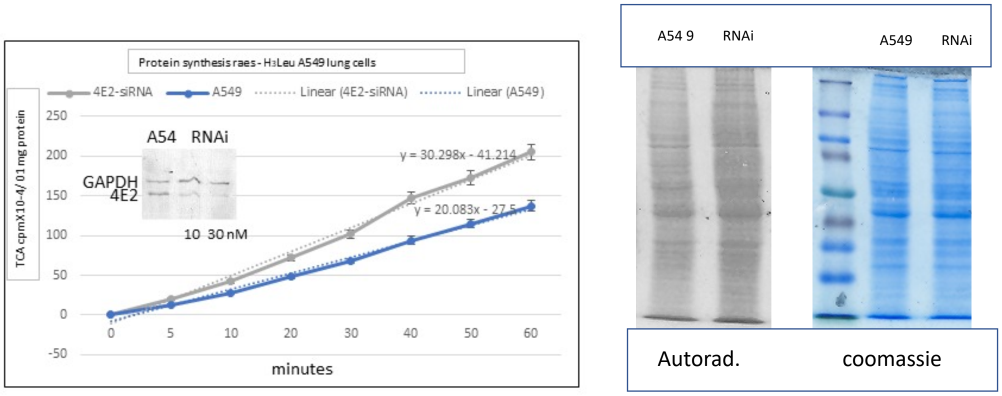


**Supplemental Figure 1. Measurement of Protein Synthesis rates in A549 cells depleted of eIF4E2**

A549 cells have been transfected with siRNA (SENSE: CGAGACAAGAAUCAGAGCAtt, Ambion/Life Technologies) for 24h. The inset shows the effective dose-depletion of eIF4E2. $10^5$ A549 cells with or without 4E2-siRNA were each plated in 12 wells of a 24-well plate with 1ml complete D-MEM and 10% FCS (duplicate samples). The next day the medium was replaced with medium containing 5µCi/ml L-[3,4,5-3H(N)]-Leucine (150 Ci/mmol - NEN), and sequential aliquots were removed at indicated intervals. After solubilization with 0.5 ml 1%SDS, 10% TCA insoluble material (0.1 mg Protein) was collected on 2.5 cm GFA filter. Following washing with 90% EtOH, the filters were air-dried and placed in scintillation vials for counting with OptiScint LLT NPE-Free Scintillation cocktail in a Beckman LS6500 counter. Note that the difference in PS rates is highly significant ($P<10^{-6}$). 10 µg of protein isolated at 1h was processed by 8%PAGE/SDS for fluorographic auradiography with PPO.

**Supplemental Figure 2. Nsp2-cells demonstrate higher cap- and HCV-dependent translation under normal and hypoxic conditions.** Expression of Renilla and Firefly luciferases from capped-Renilla-HCV-IRES-FF Luciferase mRNA in control and Nsp2 cells grown under normal (white bars) or hypoxic (grey bars) conditions. Graphs represents mean of Renilla units (expressed from the capped mRNA) **(left panel),** and Firefly (expressed from the HCV IRES mRNA) **(right panel)** luciferase signals (±SD, n=3).

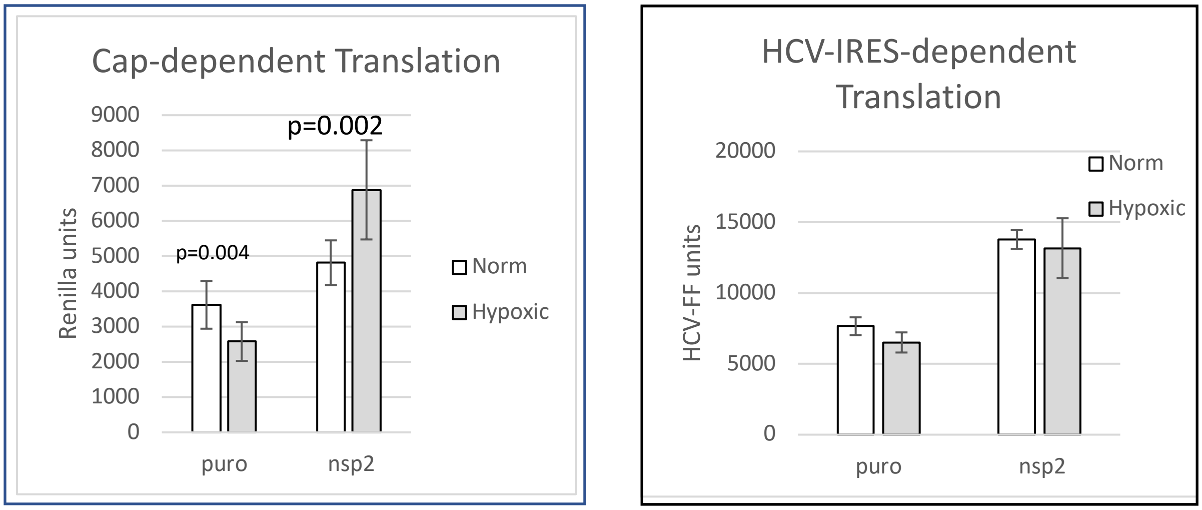


**Supplemental Figure 3 – Repeat of VEGF WB**

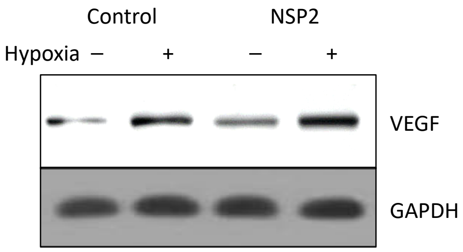
